# ATP signaling in the integrative neural center of *Aplysia californica*

**DOI:** 10.1101/2020.11.20.392001

**Authors:** János Györi, Andrea B. Kohn, Daria Y. Romanova, Leonid L. Moroz

**Author notes:** equal contribution (János Györi, Andrea B. Kohn, Leonid L. Moroz).

## Abstract

ATP and its ionotropic P2X receptors are components of one of the most ancient signaling systems. However, little is known about the distribution and function of purinergic transmission in invertebrates. Here, we cloned, expressed, and pharmacologically characterized P2X receptors in the sea slug *Aplysia californica* – the prominent model in cellular and system neuroscience. These functional P2X receptors were successfully expressed in *Xenopus* oocytes and displayed activation by ATP (EC_50_=306 μM) with two-phased kinetics as well as Na^+^-dependence. The ATP analog, Bz-ATP, was a less effective agonist (~20%) than ATP, and PPADS was a more potent inhibitor of the P2X receptors than the suramin. We showed that P2X receptors are uniquely expressed within *Aplysia*’s cerebral bioenergetic center (also known as F-cluster). Using RNA-seq, we found that the F-cluster contains more than a dozen unique secretory peptides, including three insulins, interleukins, and potential toxins, as well as ecdysone-type receptors and a district subset of ion channels. This structure is one of the most prominent integrative centers in the entire CNS and remarkably different from the morphologically similar neurosecretory center (bag cluster) involved in egg-laying behavior. Using RNA-seq, we also characterized the expression of P2X receptors across more than a dozen *Aplysia* peripheral tissues and developmental stages. We showed that P2X receptors are predominantly expressed in chemosensory structures and during early cleavage stages. The localization and pharmacology of P2X receptors in *Aplysia* highlight the evolutionary conservation of bioenergetic sensors and chemosensory purinergic transmission across animals. This study also provides a foundation to decipher homeostatic mechanisms in development and neuroendocrine systems.

**Graphical Abstract:** We show that ATP and its ligand-gated P2X receptors are essential signaling components within both the chemosensory systems and the unique integrative neurosecretory center, present in the CNS of the sea slug *Aplysia* – a prominent model in neuroscience. Expression and pharmacology of P2X receptors in *Aplysia* confirms the preservation of evolutionary conserved bioenergetic sensors across animals and provide new tools to decipher homeostatic mechanisms in neuro-endocrine systems in general.

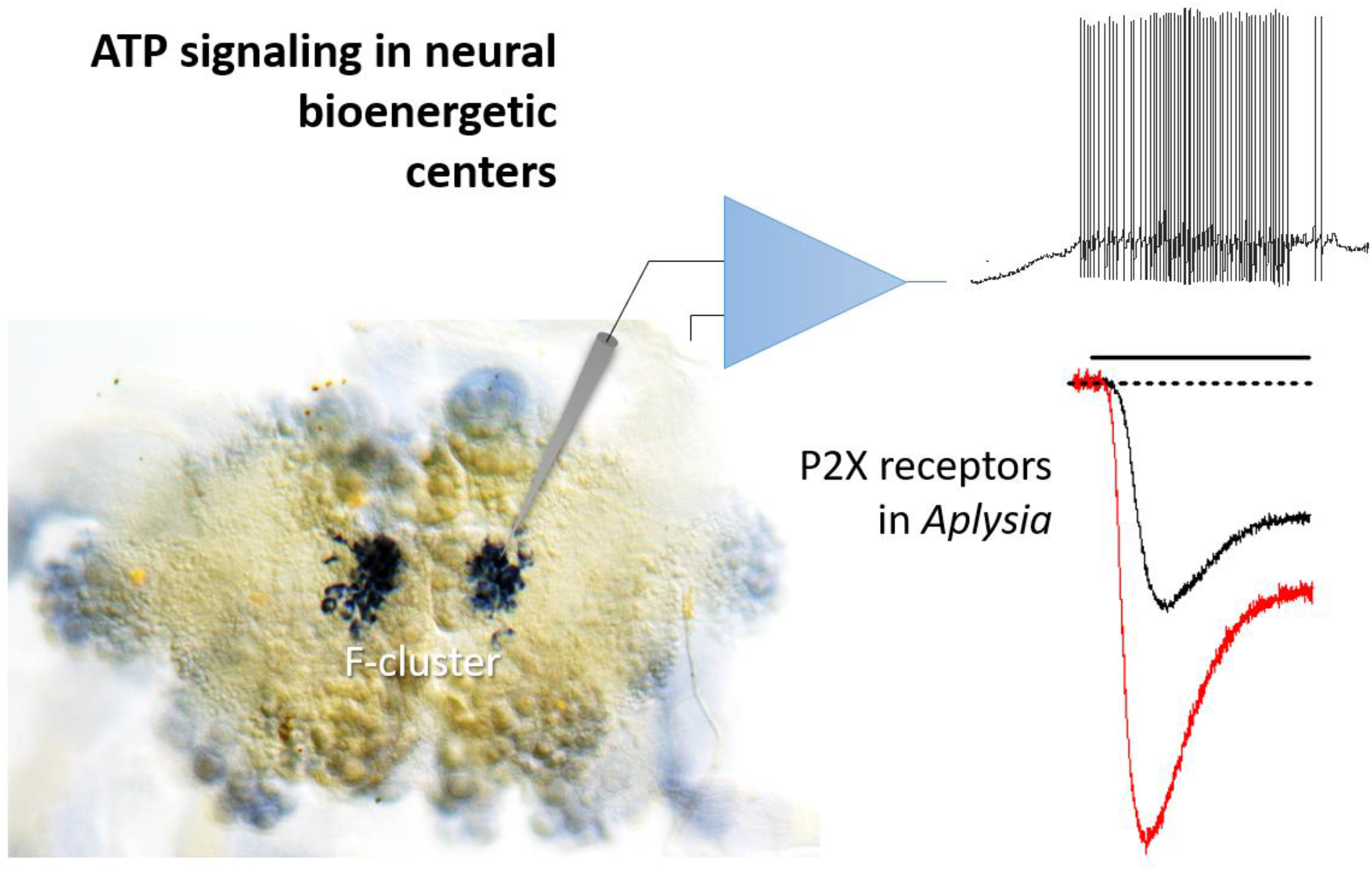

## Introduction

In addition to being the critical energy storage for every cell, ATP acts as one of the most ancient intracellular and intercellular signal molecules (1–3). The possible involvement of ATP in signaling mechanisms was initiated in the 1920s by Drury and Szent-Gyorgyi (4); and then in the 1950s by Holtons (5–7), leading to the concept of purinergic transmission in the 1970s by Burnstock (3,8). Eventually, in 1983, rapid ATP-gated ion currents were discovered in neurons (9,10) and muscles (11), and specific subtypes of the ligand-gated P2X receptors were identified in the 1990s (12–15). Finally, the 3D structure of P2X receptors has been revealed in 2009-2012 (16, 17). These are distinctive trimeric ligand-gated channels showing the common architecture with acid-sensing ion channels but unrelated in their respective amino acid sequences (18).

Comparative studies established that P2X-type receptors are broadly distributed across many eukaryotic lineages (1,19–21), including the majority of Metazoa (2,3). Across all domains of life, ATP can operate as the *bona fide* ancient signal molecule (volume transmitter) associated with adaptive reactions to injury and damage (2,3,22).

Nevertheless, the recruitment of P2X receptors into different signaling mechanisms and tissues occurred relatively randomly. In selected evolutionary lineages, P2X receptors were secondarily lost. The list includes higher plants (3), some basal bilaterians such as acoels (22), arthropods and nematodes. For example, *Drosophila* and *C. elegans* genomes have no P2X receptors, but other ecdysozoans such as *Daphnia*, the shrimp *Litopenaeus*, the tick *Boophilus* (23), and tardigrades (24) contain one receptor. Lophotrochosoans or Spiralia, including flatworms, also have one type of P2X receptor with shared pharmacological properties to mammals (25). However, practically nothing is known about the functional roles of P2X receptors in the CNS and peripheral tissues of invertebrates and molluscs, in particular (3). Mollusca is one of the most diverse animal phylum in terms of its morphological and biochemical adaptations.

The release of ATP from the central ganglia of the pond snail, *Lymnaea stagnalis* was demonstrated (26), and, subsequently, P2X receptors were identified in this species with widespread expression across the CNS (27) but unknown function(s).

Here, we show that ATP and its ligand-gated P2X receptors are essential signaling components within chemosensory structures and the unique integrative neurosecretory center present in the CNS of the sea slug *Aplysia -* an important model for neuroscience (28,29). Expression and pharmacology of P2X receptors in *Aplysia* confirms the preservation of evolutionary conserved bioenergetic reporter-sensor systems across animals and provides new tools to decipher homeostatic mechanisms in neuroendocrine systems and development.

## Results

### Identity, phylogeny, tissue-specific expression, and quantification of Aplysia P2X (AcP2X) receptors

We identified and cloned a single P2X receptor with two splice forms (GenBank#: NP_001191558.1, NP_001191559.1), which shared 92% identity (Supporting information). The predicted structure of the *Aplysia* P2X receptor reveals all major evolutionary conservative sites and posttranslational modifications (Supporting information, **Fig. 1S**), which are similar to its homolog in another gastropod, *Lymnaea* (27). The genomic organization of the P2X receptors confirmed the overall evolutionary conservation of exons and intron-exon boundaries*. Aplysia* P2X receptor exons are similar in number and length to other vertebrate P2X4 receptors, but it is not true in some other invertebrates (Supporting information, **Fig. 2S**).

**Fig. 1*A*** shows the phylogenetic relationships among P2X receptors with distinct events of gene duplications in the lineages leading to humans, zebrafishes, hemichordates, echinoderms, and basally-branched metazoans such as sponges, placozoans, and cnidarians. In contrast, representatives of molluscs (including *Aplysia*), annelids, parasitic flatworms (*Schistosoma*) seem to have a single copy of P2X-encoded genes, which often form distinct phyletic clusters within a respective phylum. This reconstruction suggests a single P2X gene in the common metazoan ancestor with independent multiplication events in selected animal lineages. It primarily occurred within vertebrates as the reflection of whole-genome duplications early in the evolution of this group. Interestingly, some bilaterians such as the acoel *Hofstenia miamia*, insects, and nematodes (30) secondarily lost P2X receptors. This mosaic-type phyletic distribution likely illustrates different system constraints for recruiting P2X receptors to novel functions or preserving ancestral molecular mechanisms of purinergic signaling.

**Figure 1.**
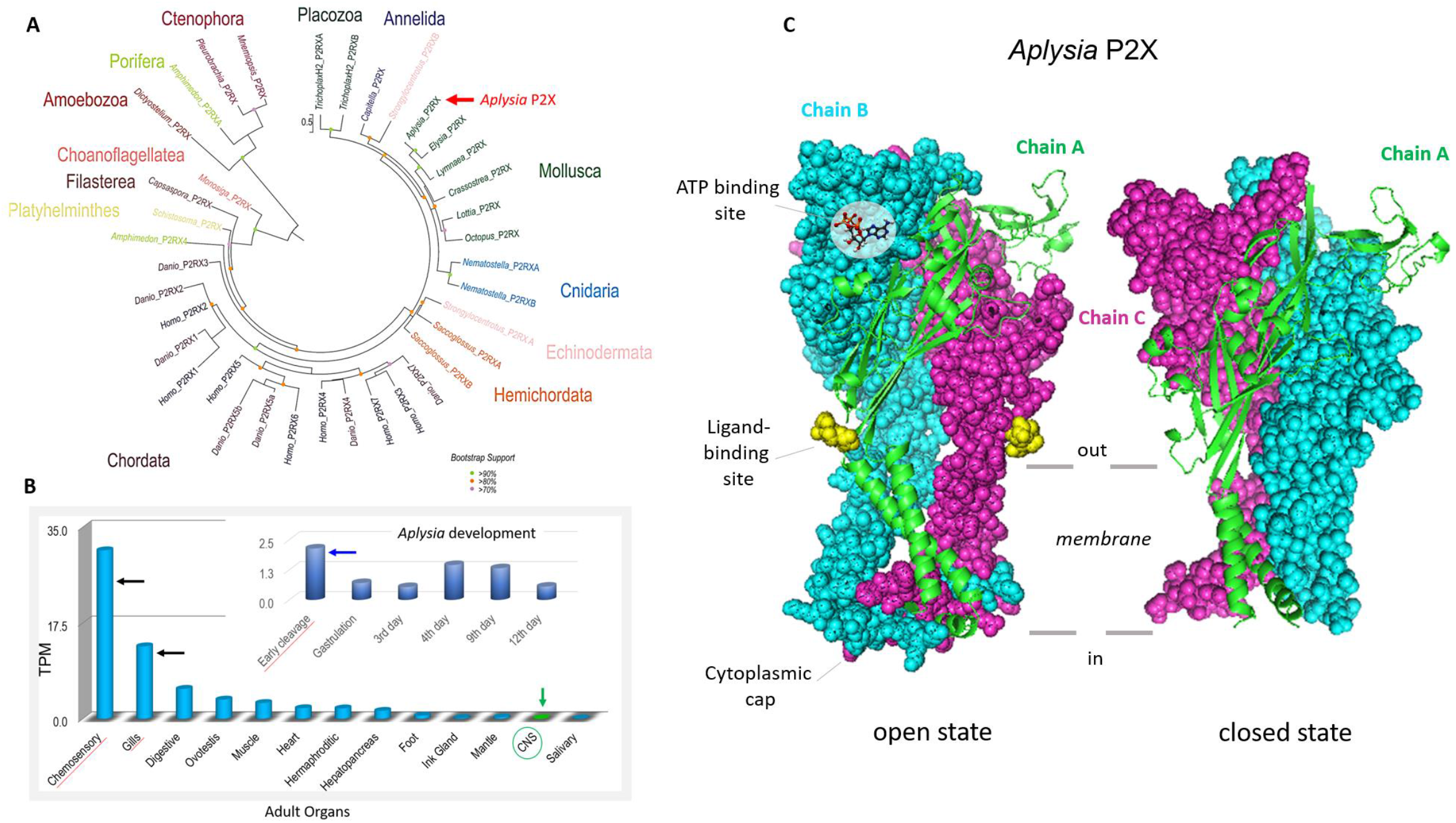
A. Phylogenetic relationships of P2X and P2X-like receptors (P2RX). ***A*.** A maximum likelihood (ML) phylogenetic tree of P2X receptors (Supporting information for sequences and accession numbers) with the best-fit model (LG+G). Bootstrap support less than 70 omitted. Phylogenetically, the P2X predicted proteins cluster by phyla. P2X-type receptors are not unique to metazoans because they are detected in unicellular green algae *Ostreococcus tauri* (20), the amoeba *Dictyostelium discoideum* (19) the unicellular eukaryote *Monosiga brevicollis* (20), as well as Capsaspora owczarzaki (21), and all these species, appear to have one P2X gene. Most of the non-bilaterians seem to have at least two P2X receptors (except for ctenophores, where only one receptor was detected). Lophotrochozoans, including the mollusc *Aplysia* and kins, appear to have one P2X receptor with different isoforms. The sea urchin and the acorn worm, *Saccoglossus*, both have at least two genes but numerous isoforms (1). Humans (13), as well as other chordates, appear to have seven unique P2X receptor genes (1). ***B*.** Quantification of the expression of P2X receptors in the CNS, peripheral tissues, and developmental stages (insert) of *Aplysia*. The RNA-seq data represented as TPM (<u>t</u>ranscript <u>p</u>er <u>m</u>illion) values (67–69). The highest expression levels were detected in chemosensory areas (mouth areas and rhinophores), gills, and early developmental stages (Supporting information, Table 1S for RNA-seq projects). ***C***. 3D modeling for P2X receptor of *A. californica* with the different states: open – A (model PDB: 5svk); closed – B (model PDB: 4dwo).

Next, we characterized the expression of P2X receptors in *Aplysia* using a broad spectrum of RNA-seq data obtained from adult and developmental stages (31) (see Supporting information, Table 1S). The highest level of *AcP2X* expression was found in the chemosensory structures (the mouth area and rhinophores (32) as well as in the gill (**Fig. 1*B***), which is also known as the chemosensory and respiratory organ. Expression of *AcP2X* receptors was also detected in the majority of peripheral organs of *Aplysia* as well as during the first cleavage stages (**Fig. 1*B***), where no neurons or specialized sensory cells exist. Thus, ATP could act as a paracrine messenger in early embryogenesis.

The freshwater pond snail *Lymnaea stagnalis* is the only know molluscan species with the biophysical characterization of P2X receptors (27). The structural organization of the *Aplysia* P2X receptor was comparable to *Lymnaea”* s P2X (**Figs. 1C, D, 2**; and **Fig. 3S** supplement) but with noticeable differences in their predicted ATP binding and other regulatory sites suggesting potentially different biophysical and pharmacological properties. These differences are also evident in the 3D models for related molluscan species (**Fig. 2** and **Fig 3S**, supplement), which suggest that marine and freshwater organisms are different in kinetic and pharmacological properties of their respective P2X receptors. This possibility was experimentally tested, as we outlined below.

**Figure 2.**
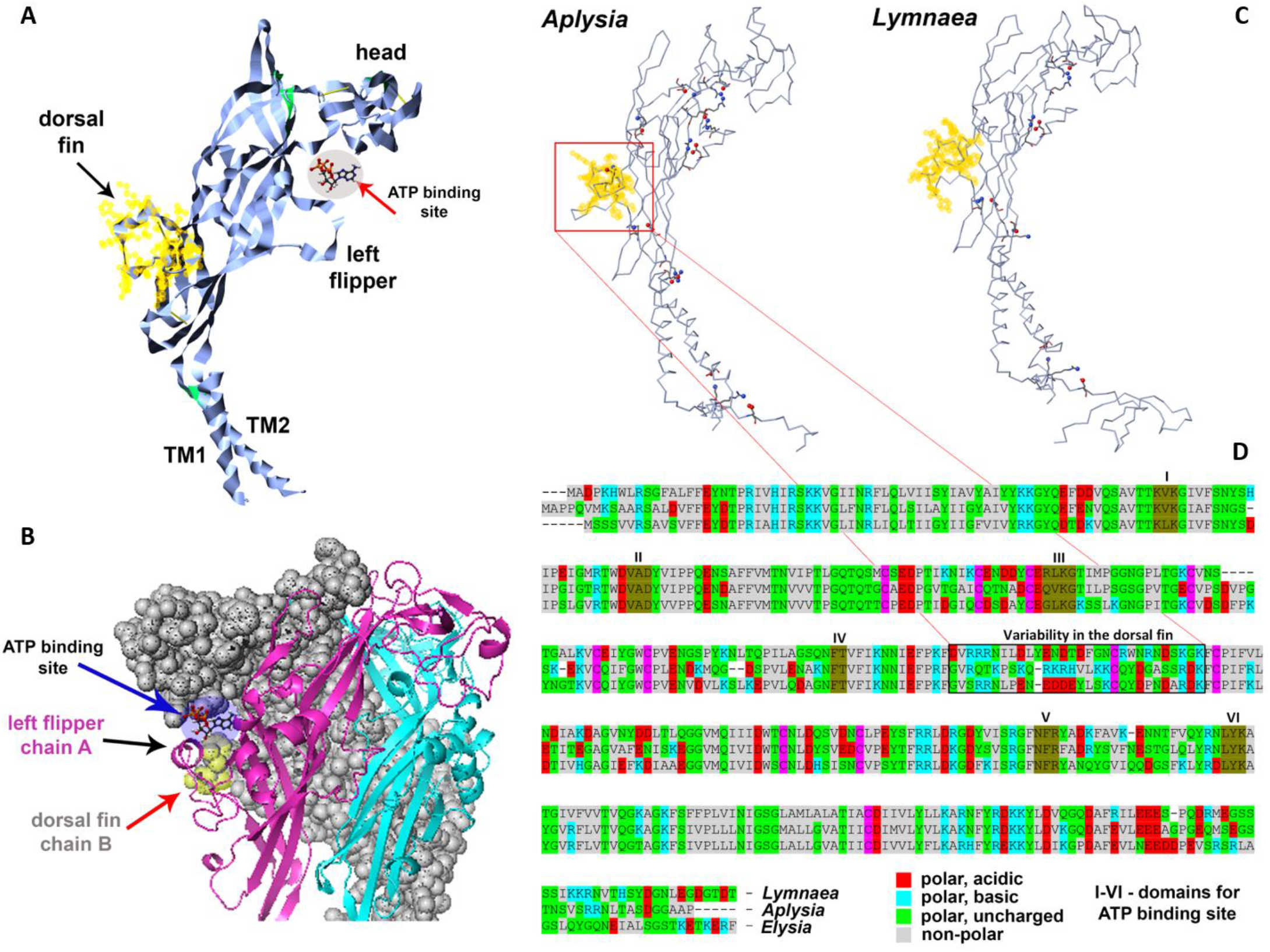
The organization of the P2X receptor in *Aplysia californica*. ***A* and *B*.** Structural features of the P2X monomer models with regions recognized for mammalian homologs in crystallography studies (16,17). TM1 and TM2 are transmembrane regions (see **Fig. 1C**). ***B***. 3D modeling of the trimeric P2X organization with suggested functional regions(16–18, 70). ‘Left flipper’ (chain A) and ‘dorsal fin’ (chain B) together with a head (chain A) of these chains form the ATP binding site (see also **Fig. 1*C***). ***C*.** Comparisons between *Aplysia* and *Lymnaea* receptors (open state) based on the difference of salt bridges (yellow – ‘dorsal fin’). ***D***. The alignment for P2X receptors in gastropod molluscs (*Aplysia californica, Lymnaea stagnalis* and *Elysia chlorotica*) with domain for the ATP binding site (brown – I-VI domains). Of note, *Aplysia* (middle in the alignment) has significantly less polar acidic and more polar basic amino acids in the ‘dorsal fin’ region [11pb:2pa] compared to other species (*Lymnaea* [7pb:6pa] and *Elysia* [5pb:8pa]), suggesting different kinetic and pharmacological properties of P2X receptors. In summary, there are 13 polar charged amino acids in the ‘dorsal fin’.

### Expression of AcP2X in Xenopus oocyte confirms the evolutionary conservation of kinetic and pharmacological parameters

ATP elicited an inward current in a concentration-dependent manner in oocytes injected with *AcP2X* (**Fig. 3*A***). EC50s were determined for both the fast-(0-1seconds) and the slow component of current with continuous application of ATP. The EC50 for the fast component was 306.0 μM with a 1.58 Hill coefficient and for the slow component 497.4 μM with a 0.97 Hill coefficient (n=5 oocytes, **Fig. 3*B***). The second application of the agonist, with a recovery time of 6 minutes, generated a 15-30% reduction in peak amplitude and is indicative of the rundown observed in other P2X receptor subunits. The response to 250 μM ATP produced a mean peak amplitude of 31.3 nA+3.8 nA and a time to a peak value of 2.76±O.2ls (n=19) with a holding membrane potential of −70 mV (**Fig. 2*C***). The ATP analog, 2’,3’-O-(4-Benzoylbenzoyl) adenosine 5’-triphosphate (Bz-ATP(33)) gave a partial response at 20% of the ATP response (n=8 oocytes, **Fig. 3*C***). There were no UTP and ADP responses within the same range of concentrations (data not shown).

**Figure 3.**
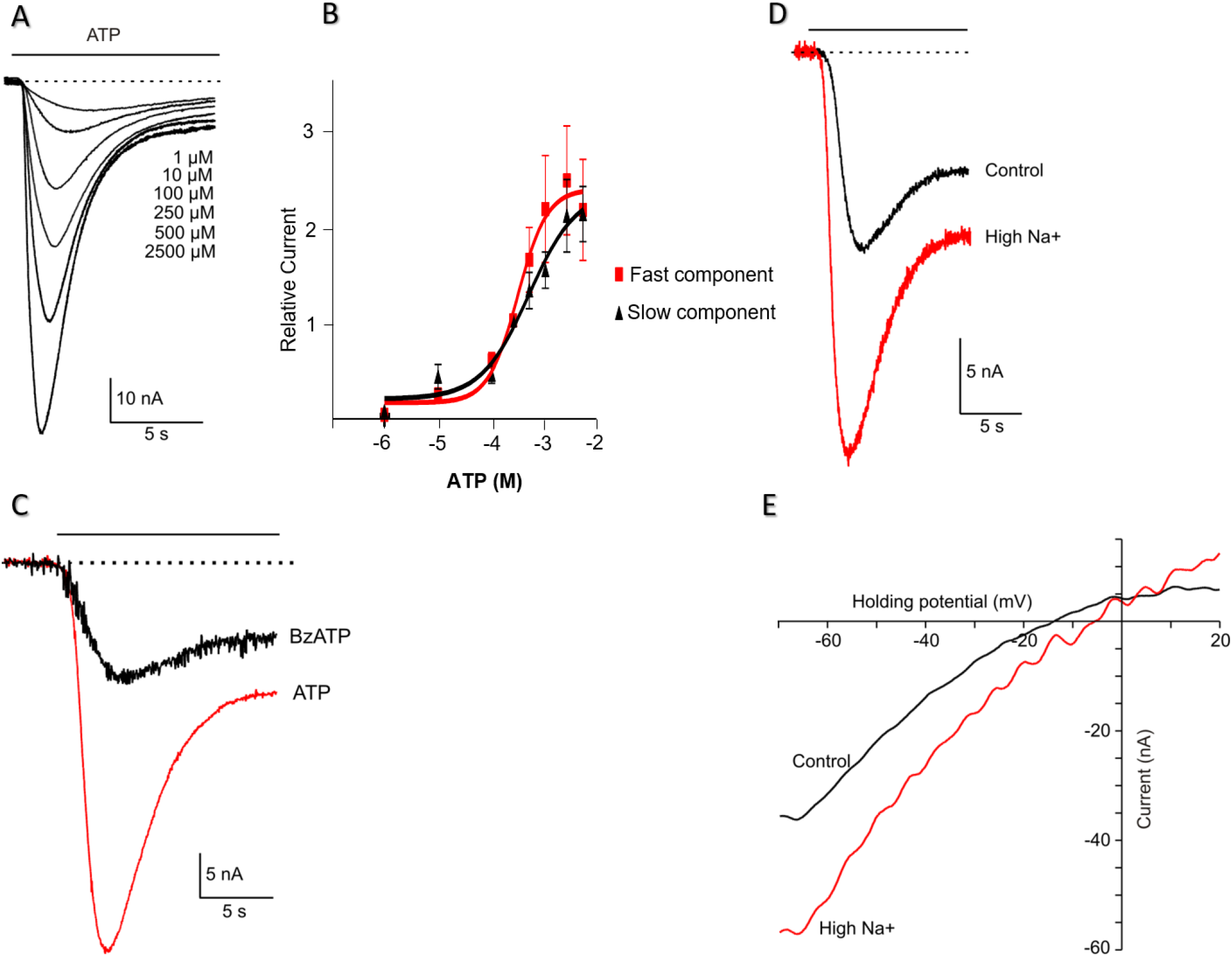
Functional expression of recombinant *Ac*P2X receptors in *Xenopus* oocytes. ***A*.** Examples of currents recorded in response to different concentrations of ATP (HP=-70 mV, agonist application indicated by the solid line). ***B***. Dose-response curves for ATP receptor activation. Mean currents were normalized to the response given by 250 μM ATP (*n=7* oocytes). Serially increasing concentrations of ATP were applied to oocytes for 15 s at 6-min intervals. Symbols represent mean±S.E. Continuous line for *ATP* represents data fitted using the equation I= I_max_/[1+(EC50/L)*nH*], where I is the actual current for a ligand concentration (L), *nH* is the Hill coefficient, and I_max_ is the maximal current (EC_50fast_ = 306.0 μM, EC_50slow_= 497.4 μM; *nH_fast_*=1.58, *nH_slow_*=0.97). ***C***. Two-electrode voltage-clamp recordings from oocytes expressing *Ac*P2X receptors. Representative inward currents recorded in response to ATP (red trace) and the 250 μM of Bz-ATP (HP=-70 mV, application indicated by the solid line). ***D***. Recordings of ATP-induced current (250 μM, ATP) in the presence of normal [Na^+^] (96mM) and with elevated extracellular Na^+^ (144 mM; red trace); HP=-70 mV. ***E***. Ramp voltage-clamp protocol from −70 mV HP to 20mV in the presence of 250μM ATP. The plots of the subtracted current (a current in the presence of ATP minus the current in the absence of ATP) against voltage during the ramp. The red trace - high [Na^+^], 144 mM. According to the Nernst equation, the reversal potential was shifted by 10.2±1.3 mV to the + direction of the *holding* potential.

The current-voltage relationship was investigated in the presence of elevated (144 mM) and low extracellular NaCl (96 mM) concentrations (n=6 oocytes, **Fig. 3*D***). A reversal potential was determined by applying a ramp protocol from −70mV to 20mV in high and normal Na^+^ with 250μM of ATP (**Fig. 3*E***). The reversal potential was 13.9mV and shifted by +10.2+1.3mV to positive holding in high sodium solution (n=6 oocytes), according to the Nernst equation.

P2X antagonist suramin (33) inhibited ATP responses in a concentration-dependent manner (**Fig. 4 *A*,*B***; 7 oocytes). Another P2X antagonist, pyridoxal-phosphate-6-azophenyl-2’,4-disulfonic acid (PPADS (33)), also inhibited the response of ATP on *Ac*P2X in a concentration-dependent manner (**Fig. 4*C*,*D***). However, the application of PPADS produced a greater block than the suramin (**Fig. 4*E***). Mean current responses to 250 μM ATP in the range of 1–250 μM PPADS generated an EC50=211.2 μM for the fast component, but the slow component could not be calculated (5-7 oocytes, **Fig. 4*D***). The second splice form of *Ac*P2Xbwas also expressed in oocytes producing currents very similar to the first isoform *Ac*P2X described above; however, it resulted in much smaller (and unstable) responses (data not shown).

**Figure 4.**
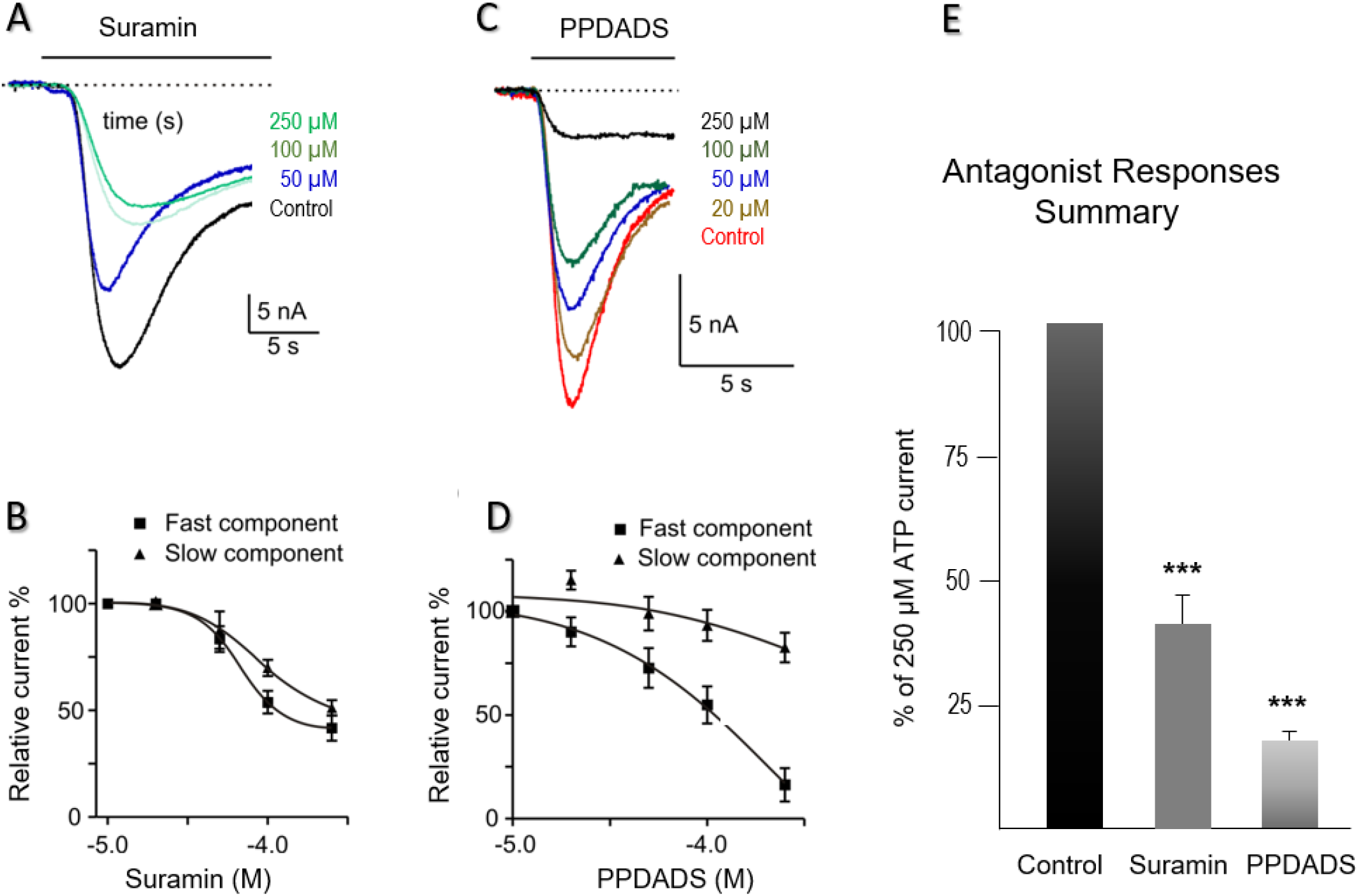
Pharmacology of *Ac*P2X receptors in *Xenopus* oocytes. ***A*.** Example of currents induced by 250 μM ATP. ATP was applied to oocytes for 15 s, in the presence of varying concentrations of suramin (HP=-70 mV). ***B***. Mean responses to 250 μM ATP in the presence of 1–250 μM suramin. There was a suramin-resistant component of the *Ac*P2X current. Symbols represent mean±S.E. ***C***. Traces recorded in response to 250μM ATP in the presence of varying concentrations of the second antagonist, PPADS (concentrations are shown in μM, and all applications are indicated by the solid lines). ***D***. Mean responses to 250μM of ATP in the presence of the PPADS (a fast component of responses - closed squares, slow component - triangles). PPADS was an effective antagonist in the range of 1–250μM. Fitting of the data using the sigmoidal dose-response curve by a continuous line, EC_50fast_=211.2. Symbols represent the mean±S.E. ***E***. The suramin proved to be a more effective blocker of the ATP-activated channels among the two antagonists tested. A chart of mean currents (% of 250 μM ATP response) in the presence of 250 μM Suramin and 250 μM PPADS. Mean currents were normalized to the response given by 250 μM of ATP. Symbols represent mean±S.E; statistically significant differences (Student’s *t*-test) from control (P<0.05) are indicated by asterisks (***) above the bars.

### The unique expression of P2X receptors in the CNS of Aplysia

Interestingly, the CNS has the overall lowest *P2X* gene expression (**Fig. 1*B***). This situation might be analogous to the recruitment of purinergic signaling in the chemosensation within the mammalian brain (34), suggesting a distinct and relatively small population of ATP-sensing cells. We tested this hypothesis.

*AcP2X* was explicitly expressed in two symmetrical subpopulations of insulin-containing neurons (**Fig. 5*A***, n=6) localized in the F-cluster in the cerebral ganglion of *Aplysia* (35,36). Each subpopulation contained about 25-30 electrically coupled cells (30-50 μm diameter, **Fig. 5*A-B***) (35). Application of 2 mM ATP to these neurons elicited a 2-5 mV depolarization, action potentials, and these effects were reversible (**Fig. 5*C***) and voltage-dependent (**Fig. 5*D***), consistent with the pharmacological properties of *Aplysia*’s P2X receptors expressed in oocytes. Neurons that were negative for *AcP2X* by *in situ* hybridization showed no response to as high as 10 mM ATP concentration.

**Figure 5.**
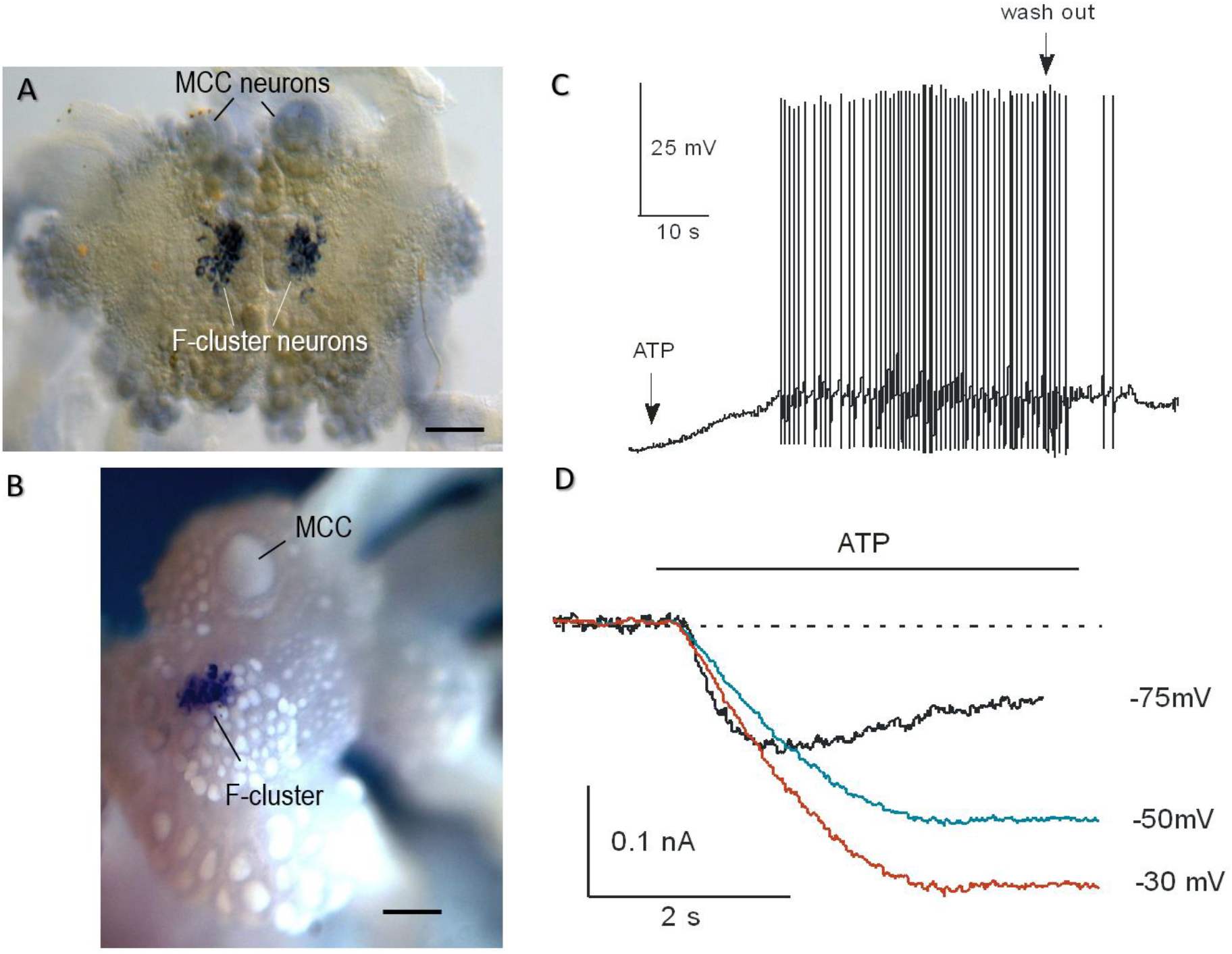
Distribution of *AcP2X* in the CNS of *Aplysia* and the effect of ATP on *Aplysia* F-cluster neurons. ***A*** and ***B***: *AcP2X*is expressed in neurons of the cerebral F-cluster (*in situ* hybridization). A pair of giant serotonergic feeding interneurons (MCC) are indicated by arrows. ***A***. The preparation embedded in a mounting media. ***B***. The cerebral ganglion was photographed in 100% ethanol. Scale: 300 μm. ***C*.** Current-clamp recording from F-cluster neurons in the intact CNS. Bath application of ATP (2.0 mM) caused an excitatory response with spiking activity (2-5 mV depolarization with a burst of the action potentials), and full recovery following washout (indicated by arrows). ***D***. Voltage-clamp recording from F-cluster neuron. Raw traces recorded in response to 2.0 mM of ATP at three holding potentials (agonist application indicated by the line).

These tests confirmed that P2X receptors in F-cluster neurosecretory cells are functional. Next, using the RNA-seq approach, we investigated this cluster’s molecular organization to get insights into biology of this cell population.

### Molecular organization of two neurosecretory centers: F-cluster vs Bag cells

We compared F-cluster molecular organization (**Fig. 6*A***) to other neurosecretory cells with a similar overall morphological organization, such as the bag cluster neurons (**Fig. 6*B***) in the abdominal ganglion (37). Neurons from both of these clusters are the most prominent secretory cells in the entire organisms (**Fig. 6*D***); both groups of cells release their secretes in neurohaemal areas along the nerves and their connective tissues (35,37) but their molecular (and perhaps systemic) functions are remarkably different. This situation is clearly reflected in their distinct transcriptome profiles (**Fig. 6**, and Supporting information **Figs 4S–7S**). There are 5159 genes differentially expressed in F-cluster (Supporting information, Table 2S).

**Figure 6.**
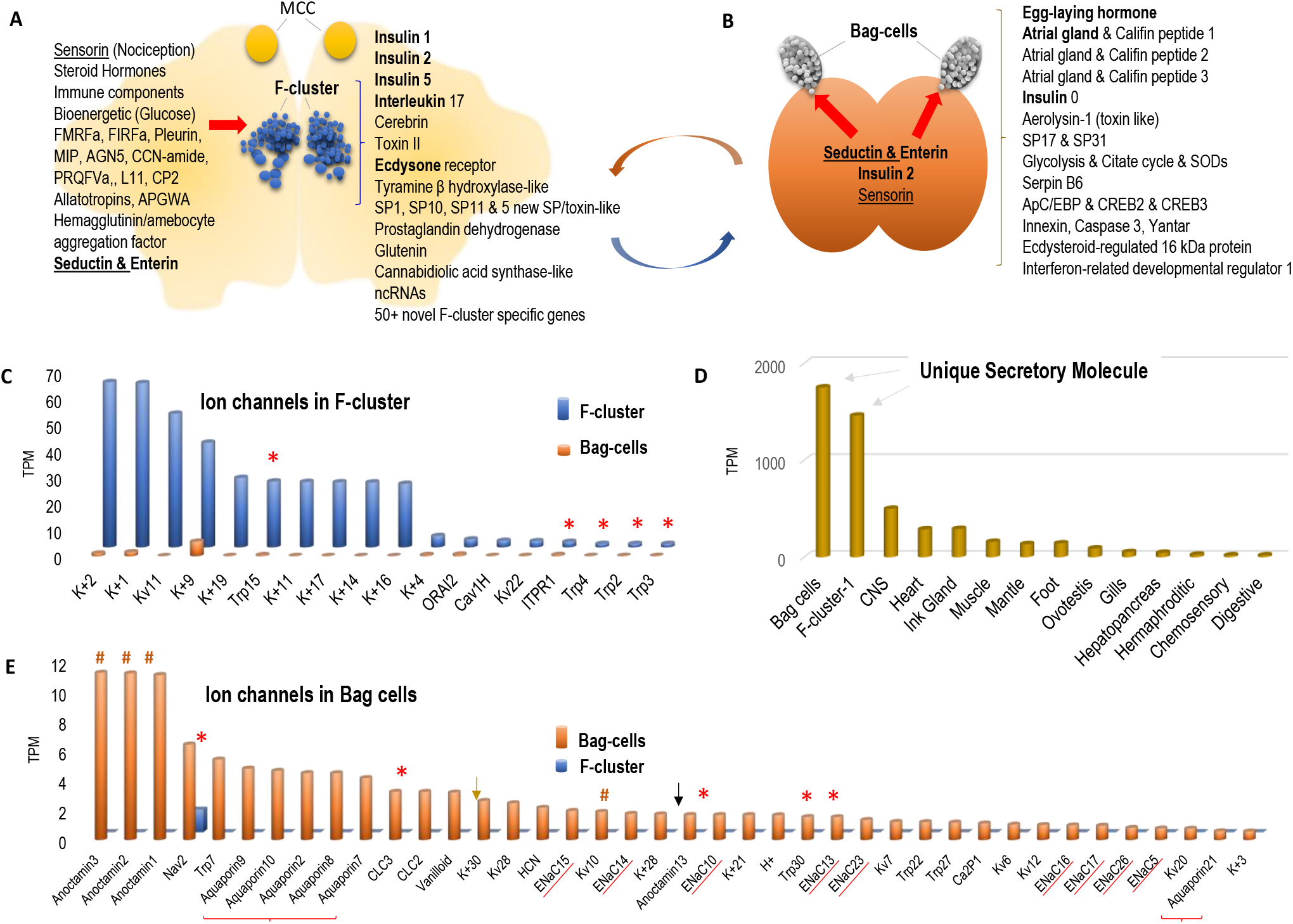
F-cluster and Bag cells are the two most prominent neurohaemal centers in *Aplysia*. ***A*.** Schematic representation of F-cluster in the cerebral ganglion (MCC-serotonergic metacerebral cells involved in feeding arousal) together with genes [on the right, e.g., insulins 1 and 2, interleukin 17, etc.] differentially expressed in this cluster (RNA-seq). Red arrow indicates predicted other peptidergic inputs [on the left] to F-cluster (see text for details). F-cluster is one of the most molecularly heterogenous neurorosecretory clusters in the CNS. ***B*.** Bag cells in the abdominal ganglion with cluster-specific peptides [on the right] expressed in these neurosecretory cells (RNA-seq). Red arrow indicates other peptidergic inputs to bag cells (see text for details). Of note, seductin, enterin, and sensorin inputs are shared (underlined) between these two neurohaemal centers, whereas insulin 2 is produced in the F-cluster. ***C*.** Both bag cells and F-cluster have the highest overall expression of an uncharacterized secretory molecule (XP_005092953.2) compared to other organs. ***D*** and ***E***. Putative ion channels differentially expressed in the F-cluster and bag cells, respectively (see Supporting information excel table:Ion channels for F-cluster and Bag cells). Asterisks indicate cluster-specific transient receptor potential (TRP) channels; # - anoctamins,– aquaporins; brown and black arrows HCN and voltage-gated proton channels, respectively. ENaC – are ligand-gated peptide channels (underlined). Other abbreviations: K+ Calcium-activated, Inwardly rectifying, Tandem pore domain potassium channels; Kv voltage-gated potassium chnnels; Nav-voltage-gated sodium channel, Ca2P two pore calcium channels. The numbers in names Do Not infer type of channel but for identification only.

The neurosecretory bag cells induce and integrate complex egg laying behaviors due to the release of several secretory peptides (38–40). Predictably, all of them were highly expressed in our RNA-seq dataset (**Fig. 6*B***, Supporting information, Table 2S), but there were 7213 genes expressed only in bag cluster, including a distinct isoform of *insulin, aerolysin* 1-toxin-like peptide, and a diverse set of ion channels (**Fig. 6*C*** and ***E***, Supporting information, Table 2S).

In contrast to bag cells, very little is known about P2X expressing F-cluster neurons. It was shown that these cerebral neurons contain insulin and have broad secretory areas with a likely non-synaptic hormone release in the connective tissues, nerves and circulatory system (36). It was suggested that these neurons may be involve in glucose control (41) and perhaps reproduction, with likely hormonal interactions to the bag cells.

Our RNA-seq data (**Fig. 6*A***, supporting information, Table 2S) revealed that F-cluster neurons expressed three distinct *insulins*, *cerberin*, and at least 4 novel secretory neuropeptides as well as toxin-like peptides. Most surprising finding in F-cluster neurons was an extremely high level of expression of *interleukin 17A*-like (a pro-inflammatory cytokine (42)) and *interferon α-inducible protein 27* (*IFI27*), originally known to be involved in innate immunity, and is later found to intervene in cell proliferation. An *Aplysia*-type ecdysone receptor was also differentially expressed in F-cluster neurons consistent with expression of many components of steroid hormone machinery in *Aplysia* (Supporting information, Table 2S). Also, and in contrast to bag cells, we found evidence of innervation of F-cluster neurons by about a dozen of neuropeptides (*FMRFa, FIRFa, Pleurin, MIP, AGN5, CCN-amide, PRQFVa, L11, CP2, Allatotropins, APGWA*) including *sensorin* from nociceptive neurons (43). In each of these cases, it is known (36,41,44,45) that these neuropeptide prohormones are expressed in different neuronal populations (our *in situ* hybridization for these transcripts, not shown), but respective precursor mRNAs are transported to distant neurites (45) and these extrasomatic RNAs are detected from synaptic regions on target neurons.

Of note, two neuropeptide inputs mediated by *seductin* (a pheromone involved in integration of reproductive behaviors (46–48) and *enterin* (myoactive factors (49–52)) are apparently shared between F-cluster and bag cells (**Fig. 6*A*, *B***). Together with the presence of insulin receptors in bag cells, it implies long-distance interactions between F-cluster neurons and bag cells.

Non-coding RNAs also showed differential expression between clusters (Supplementary Table 2S).

## Discussion

As the central bioenergetic currency, the intracellular concentrations of ATP reach 1-10 mM with multiple mechanisms of its extracellular release across all domains of life (2). In the molluscan CNS, the baseline level of ATP release can be increased following depolarization and serotonin application, suggesting that ATP can act as an endogenous fast neurotransmitter (26). The presence of ionotropic P2X ATP-gated cationic channels in peripheral chemosensory structures (together with profound serotonergic innervations of these structures (53)) and the CNS (**Fig. 1*B***) further support this hypothesis. However, we also reported *AcP2X* receptor expressions early in development, suggesting that ATP might be a paracrine signal molecule controlling cleavage and differentiation.

The purinergic sensory transmission is widespread in mammals (34) and might have deep evolutionary roots (2,3,22). Mammalian P2X receptors (13) are comparable to their homologs in *Aplysia* based on sensitivity to ATP and kinetic parameters. The overall kinetic and pharmacological parameters of *Aplysia* P2X receptors are also similar to those described both in the closely related *Lymnaea* (27) and distantly related *Schistosoma* (25). However, the *Lymnaea* P2X receptor showed much higher sensitivity (EC50 is in μmolar range) to ATP than *Aplysia*, consistent with structural differences of P2X receptors across species (**Figs. 1–2**). Suramin and PPADS both inhibited the ATP evoked responses in other species (23–25, 27), but in *Aplysia*, it occurred in a narrower range (10-250 μM) than in *Lymnaea* (0.1 μM-250 μM).

Ecdysozoan P2X receptors are relatively diverse. In contrast to *Aplysia*, the tick P2X receptor displayed a very slow current kinetics and little desensitization during ATP application (23). The tardigrade P2X receptor (24) had a relatively low sensitivity for ATP (EC50 ~44.5 μM), but fast activation and desensitization kinetics - similar to *Aplysia*.

Thus, *Aplysia* P2X receptors exhibit a distinct phenotype, having a moderate ATP sensitivity (compared to the freshwater *Lymnaea*) but faster kinetics than some ecdysozoans. These “hybrid features” might be related to the marine ecology of *Aplysia* with a wider range of environmental changes.

Interestingly, the abundance of P2X receptors in the *Aplysia* chemosensory systems (such as mouth areas, rhinophores, and gills) correlates with the expression of nitric oxide synthase (54), suggesting interactions of these afferent pathways in the control of feeding and respiration. *Aplysia* might also detect environmental ATP from bacterial and algal (food) sources (2) as in some other studied marine species, including lobsters (55).

Within the CNS of *Aplysia*, P2X receptors are expressed in the distinct cluster of insulin-containing neurons (35,36), likely associated with the systemic control of growth and, subsequently, reproduction. The potential functional roles of many differentially expressed genes in F-cluster could be determined in the future but their involvements in control of multiple behaviors, immunity is evidenced by a broad array of respective genes uniquely or differentially expressed in F-cluster (Supporting information, Table 2S). This is the reason to view this cluster as one of the top-level integrative centers in the animal with the broadest spectrum of secretory peptides and multiple receptors, including P2X receptors.

The release of the *Aplysia* insulin can decrease the level of glucose in the hemolymph (36). Moreover, F-cluster neurosecretory cells are electrically coupled (35), which help synchronize their discharges and, eventually, insulin secretion. It is known that ATP can also be released from gap junction (innexins) and during synaptic exocytosis (2). Thus, we can propose that P2X-induced neuronal depolarization of insulin-containing neurons provide positive purinergic feedback sustaining the excitability and secretory activity of this multifunctional integrative center in *Aplysia* and related gastropods.

In summary, the observed diversity of specific transcripts controlling pain, injury, toxins, immunity feeding, reproduction and energetics, together with ATP-sensitivity suggests that F-cluster can represent a neurosecretory integrative center, controlling (at least in part by purinergic signaling) a broad majority of the animal’s behavior and functions.

## Experimental procedures

### Molecular analyses of Aplysia AcP2X

*Aplysia californica* (60-100 g) were obtained from the National Resource for *Aplysia* at the University of Miami (Supporting information for details). The original sequences were generated using RNA-seq profiling (41,44,45,56). Details for RNA extraction and cDNA library construction have been described (41,44,56,57) and provided in Supporting information. We used the same protocols for whole-mount *in situ* hybridization as reported elsewhere (58,59) with a specific probe for the validated *Ac*P2X (Supporting information). Expression of *AcP2X* was performed in eight experimental and two control preparations of the CNS; additional controls were reported elsewhere (44,59). Control *in situ* hybridization experiments with full length ‘sense’ probes revealed no specific or selective staining under identical labeling protocols. Images were acquired with a Nikon Coolpix4500 digital camera mounted on an upright Nikon Optiphot-2 microscope.

Expression levels of transcripts were calculated using the normalization method for RNA-seq - Transcripts Per Million (TPM) (60). Mapping was performed in the STAR (2.3.0)/feature Counts analysis with the values obtained from the Bowtie2/Tophat pipeline (61). The mapped reads were summarized and counted within the R statistical programming language. Supporting information, Table 1S contains a list RNA-seq projects and their corresponding SRA number.

### Electrophysiology

#### Oocytes recordings

RNA preparation for oocyte injections and oocyte maintenance are found in Supporting data. The oocyte recording bath was in ND96 medium (96 mM NaCl, 2 mM KCl, 1 mM MgCl_2_, 1.8 mM CaCl_2_, and 5 mM HEPES, pH=7.4) with the 1.8mM CaCl_2_ being replaced by 1.8 mM BaCl_2_. Whole-oocyte currents were recorded by two-electrode voltage clamp (GeneClamp500B,Axon Instruments, Foster City,CA,USA) using microelectrodes made of borosilicate glass (WPI,USA) with a resistance of 0.5–1 MΩ when filled with 2.5 M KCl. Currents were filtered at 2 kHz and digitally sampled at 5 kHz with a Digidata 1320B Interface (Axon Instruments,CA). Recording and data analysis were performed using pCLAMP software version 8.2 (Axon Instruments). For data acquisition and clamp protocols, the amplifiers were connected via a Digidata 1320B AD/DA converter (Axon,USA) to an AMD PC with pClamp 8.2 voltage-clamp software (Axon,USA). Unfiltered signals were sampled at 10 kHz and stored digitally.

Data are presented as mean+S.E. using Student’s paired *t*-test. Concentration-response data were fitted to the equation I=I_max_/[1+(EC_50_/L)^*nH*^], where I is the actual current for a ligand concentration (L), *n_H_* is the Hill coefficient, Imax is the maximal current and EC50 is the concentration of agonist evoking 50% the maximum response. To compute the reversal potential for sodium the Nernst equation used; Vj=(RT)/(zF)ln(c1/c2) where R is the gas constant 1.98 calK^-1^mol^-1^, F is the Faraday constant 96,840 C/mol, T is the temperature in ^o^K and z is the valence of the ion.

#### *In situ* recordings

Voltage- and current-clamp experiments were carried out on identified F-cluster neurons in intact nervous systems of *Aplysia* (35). ~0.5 mL bath was perfused with solutions using a gravity-feed system and a peristaltic pump, and solution exchanges were performed by VC-6 six-channel valve controller (Warner Inst., USA). Conventional two-electrode (3-10 MΩ) voltage-clamp techniques (Axoclamp2B, TEVC mode) were employed to measure agonist-activated currents as reported(62) at room temperature(20±2°C). To characterize membrane and action potentials, we used a bridge mode of Axoclamp2B with borosilicate microelectrodes (tip resistance: 10-18 MΩ, with 0.5 M KCl, 2 M K-Acetate, and 5 mM HEPES, pH=7.2).

#### RNA-seq

The isolation of specific cells, RNA extraction and preparation of cDNA for RNA-seq analysis as well as transcriptome annotation has been performed using the same methods as reported elsewhere (41,44,56,57).

#### Protein modeling

The reconstruction of 3D-structures of the P2X receptor from *Aplysia californica* (NP_001191558.1, GenBank, NCBI) and *Lymnaea stagnalis* (AFV69113.1, Genbank, NCBI) was based on pdb ID: 5svk (open state) and 4dwo (closed state) modeling (63). Alternative models of the same P2X receptors were generated using PyMol (The PyMol Molecular Graphics System, Version 1.8.6.0 Schrödinger, LLC) and Phyre2 software (63–66).

## Data availability

All data are contained within the article as well in supporting information.

## Acknowledgments

We thank Dr. T.Ha and E. Bobkova for help with cloning and *in situ* hybridization.

## Author contributions

All authors had access to the data in the study and take responsibility for the integrity of the data and the accuracy of the data analysis. Research design, acquisition of data: all authors. Molecular data, expression and RNA-seq: A.B.K., L.L.M.; Protein modeling and analysis: D.R. Pharmacological tests: J.G.; Analysis and interpretation: all authors: Manuscript writing: L.L.M. Funding: L.L.M.

## Funding and additional information

This work was supported by the Human Frontiers Science Program (RGP0060/2017) and National Science Foundation (1146575,1557923,1548121,1645219) grants to L.L.M. Research reported in this publication was supported by the National Institute of Neurological Disorders and Stroke of the National Institutes of Health under Award Number R01NS114491 (to L.L.M.). The content is solely the responsibility of the authors and does not necessarily represent the official views of the National Institutes of Health.

## Supplementary Information 1

### Supplementary Figures

**Supplementary Figures**

**Figure 1S.**
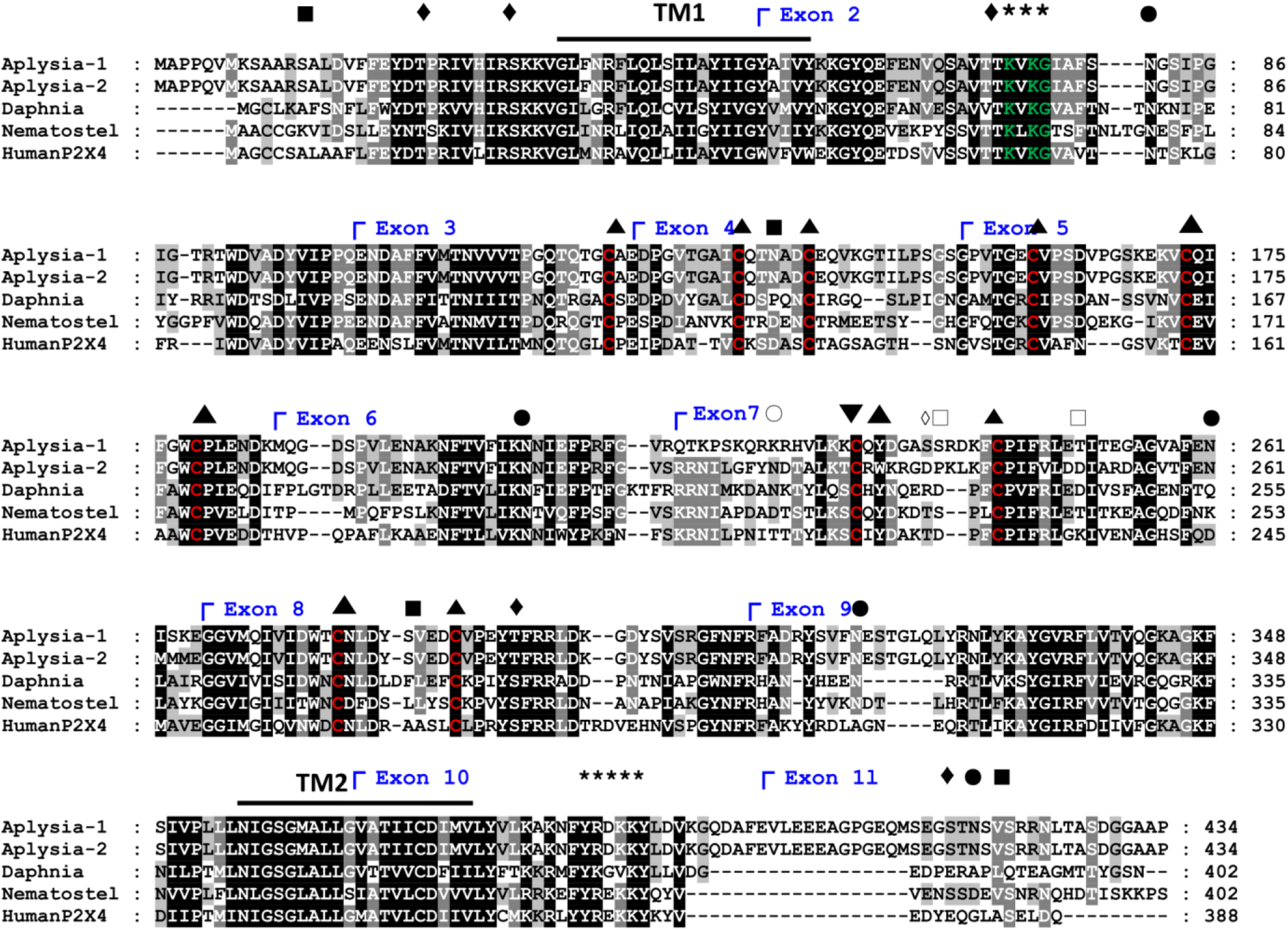
Alignment of predicted amino acid sequences for the P2X receptor subfamily. The predicted amino acid sequences for *Ac*P2X and *Ac*P2Xb were aligned with P2X4 like receptors, see materials and methods section 2.3 (the main text and above). The names and accession numbers for sequences used are: *Aplysia-1 Ac*P2X, NP_001191558.1; *Aplysia-2 Ac*P2Xb, NP_001191559.1; *Daphnia pulex*, EFX89098.1; *Nematostella vectensis*, XP_032239188.1; *Homo sapiens*, HumanP2X4, NP_002551.2. The exon/intron boundaries, as well as protein motifs, are based on the genomic or predicted protein sequence of *Ac*P2X unless otherwise noted. The key for marked critical motifs or amino acid residues is as follows: **-** Transmembrane region; • N-glycosylation site; ◆ PKC phosphorylation site; ■ CK2 phosphorylation site; ◇□○ Present only in *Ac*P2X with the same designation as filled in symbols; ▼ Present only in *Ac*P2X_b_ (PKC phosphorylation site); ▲ Conserved cysteines (red); *** Predicted ATP binding site (green); ***** Predicted trafficking motif; ┌ Exon/Intron Boundaries (blue)

**Figure 2S.**
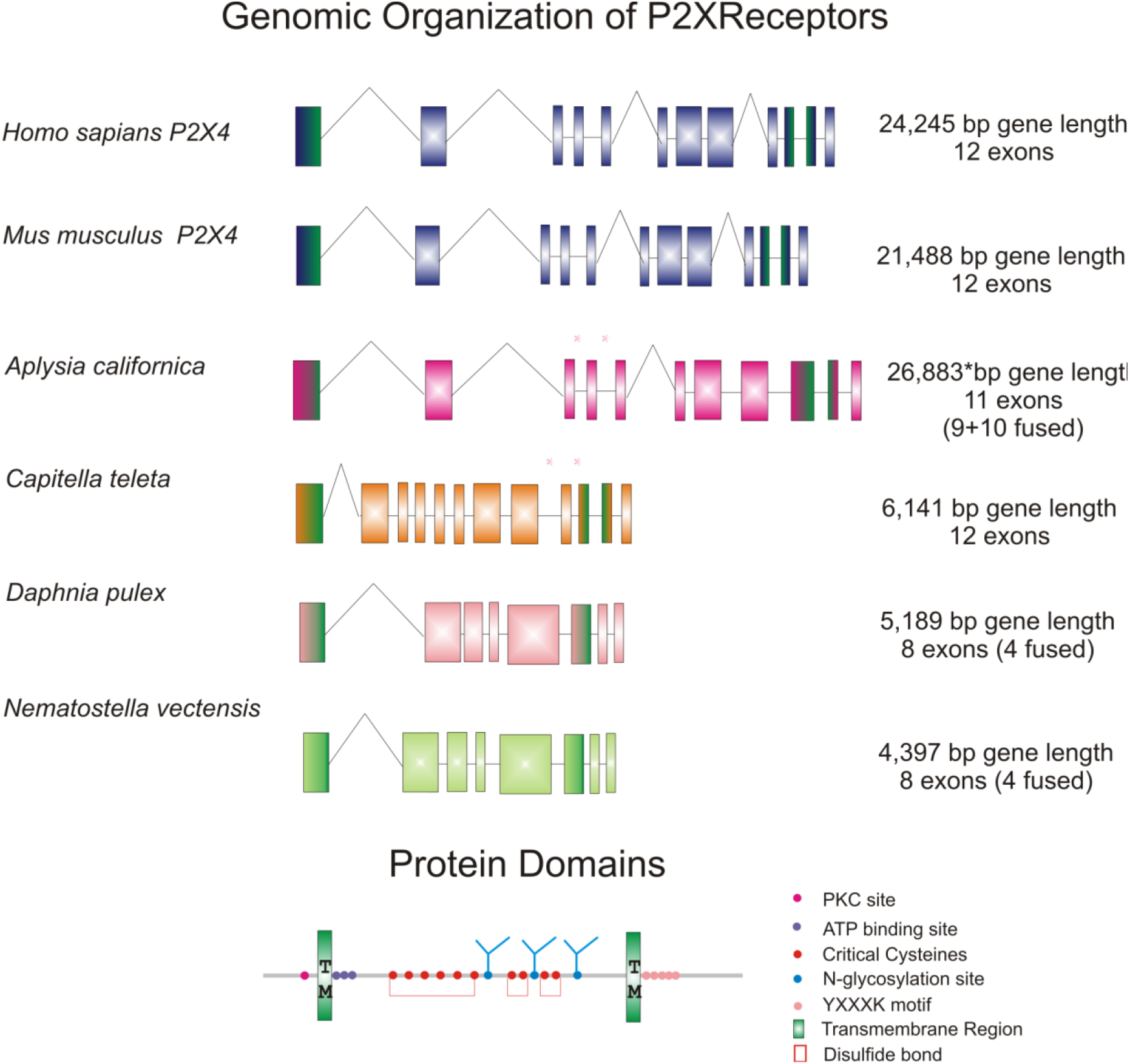
Genomic organization of P2X4 receptors. The genomic organization of selected P2X genes (same sequences and accession numbers used as in **Fig. 1S**, see also supplement 2) Human P2X4 and mouse P2X4 have 12 coding exons. *Aplysia* P2X receptor genes have 11 coding exons in which the 9 and 10 exons compared to vertebrates are fused, still generating a conserved exon/intron boundary. The asterisks indicate two contigs in the *Aplysia* genome assembly. All the exon/introns boundaries conserved and have compatible gene size. The two other invertebrate P2X-like receptor sequences from *Daphnia* and *Nematostella* only have 8 coding exons with 4 exons being fused, however, the exon/intron boundaries of those present in their genomes are still conserved. The exons are color-coded with the appropriate protein domain as well as critical amino acids being indicated.

**Fig. 3S.**
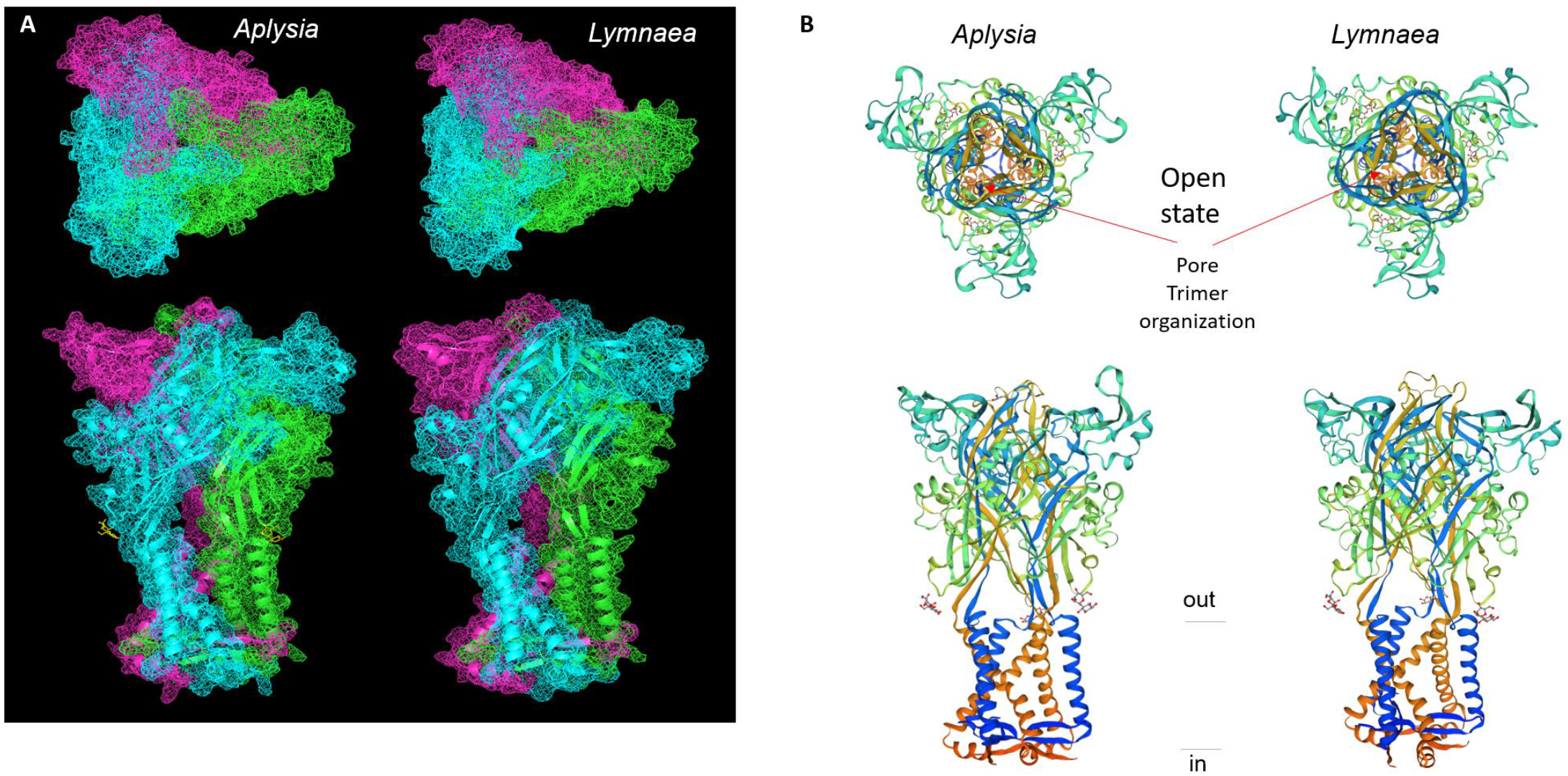
The hypothetical 3D-structure of P2X receptors for *Aplysia californica* and *Lymnaea stagnalis*. **A and B.** 3D modeling for P2X receptors of *A. californica* and *L. stagnalis* with an open state (model PDB: 5svk), see details in (66). Alternative models of P2X receptors were generated using PyMol (The PyMol Molecular Graphics System, Version 1.8.6.0 Schrödinger, LLC) and Phyre2 software (63–66).

**Fig. 4S.**
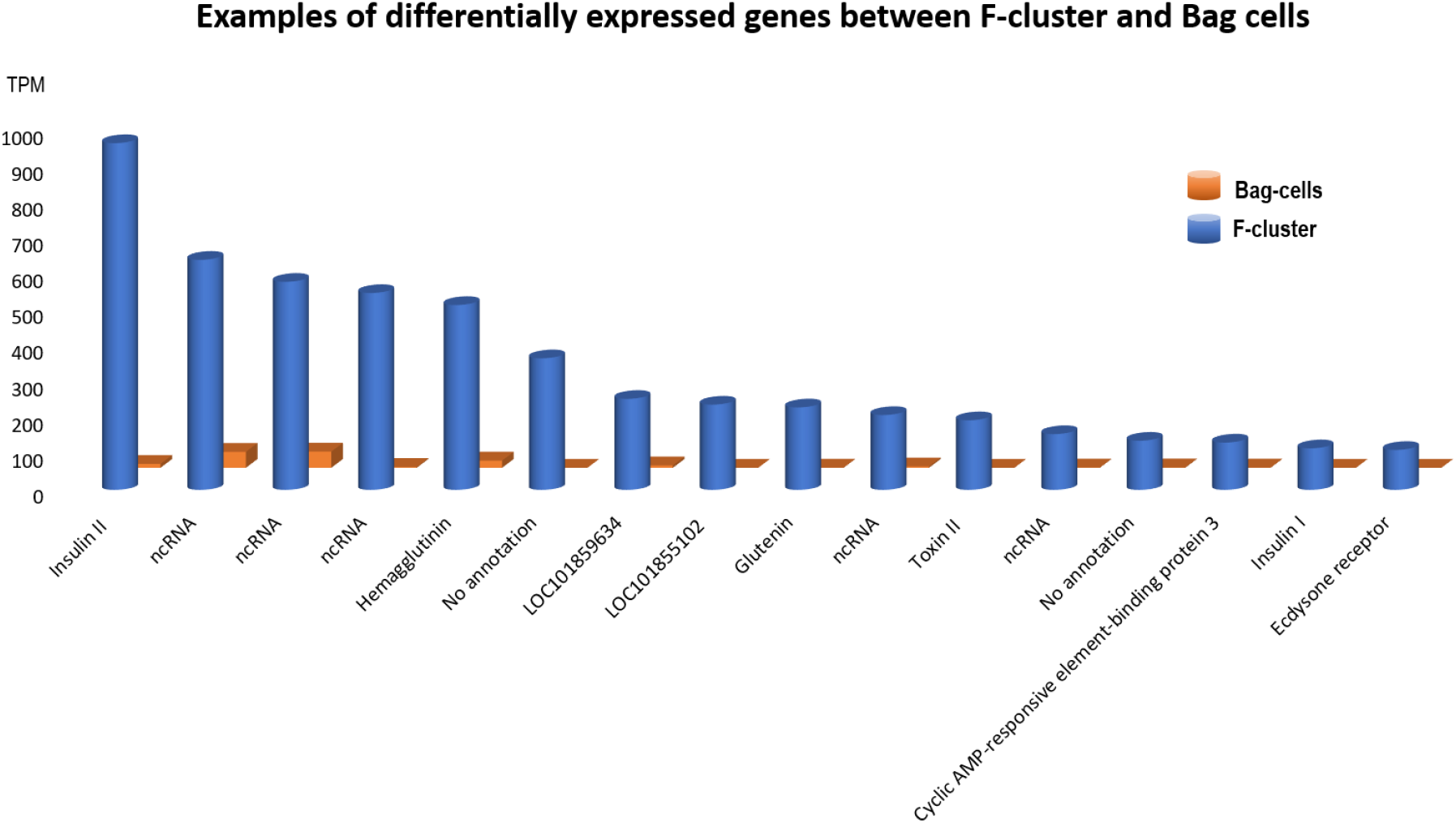
Examples of differentially expressed genes between F-cluster and Bag cells (RNA-seq). TPM - transcripts per million. See text for details.

**Fig. 5S.**
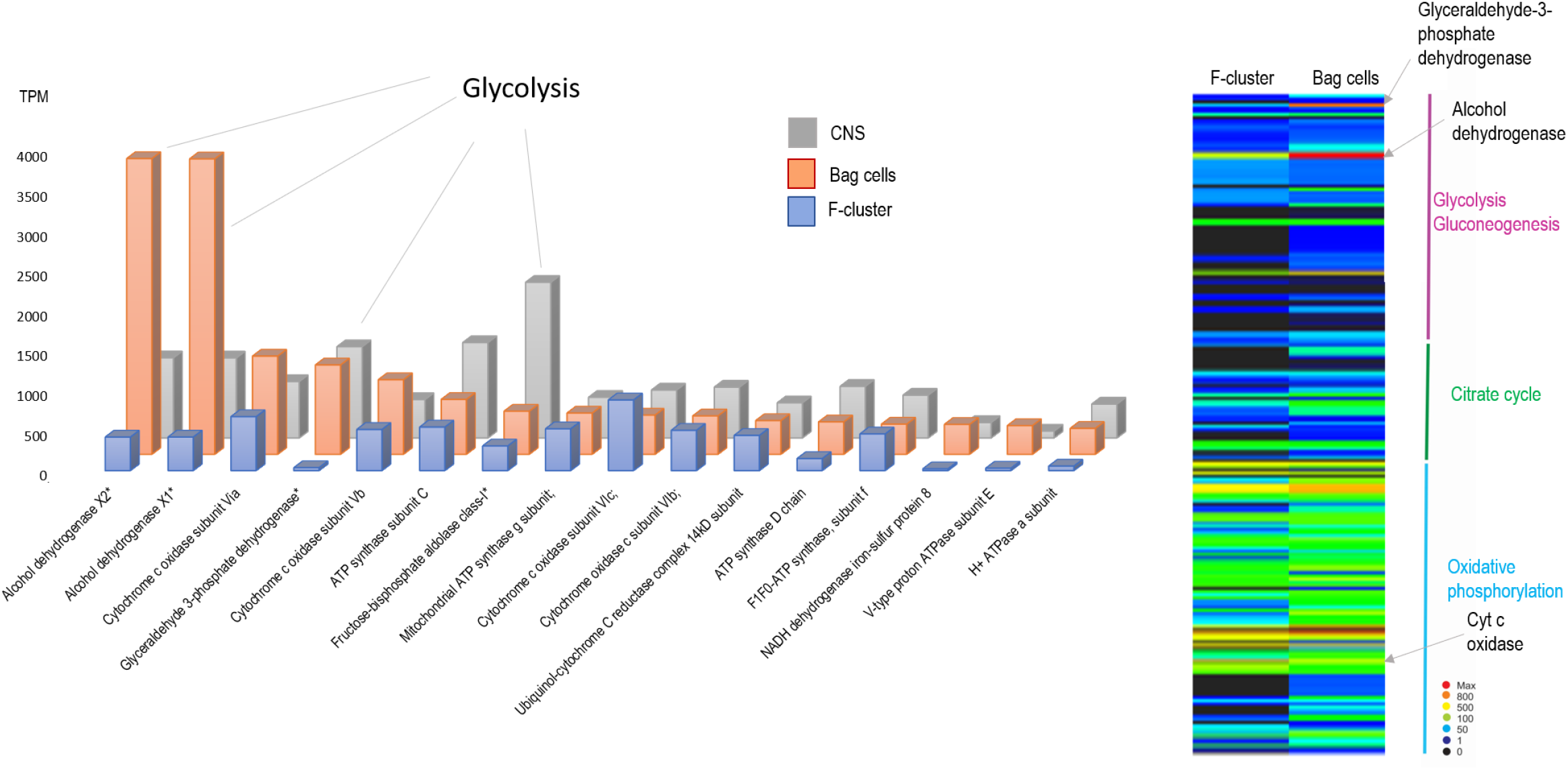
Examples of differentially expressed genes associated with bioenergetics. RNA-seq data for F-cluster, Bag cells and the entire central nervous system (CNS). TPM - transcripts per million. See text for details.

**Fig. 6S.**
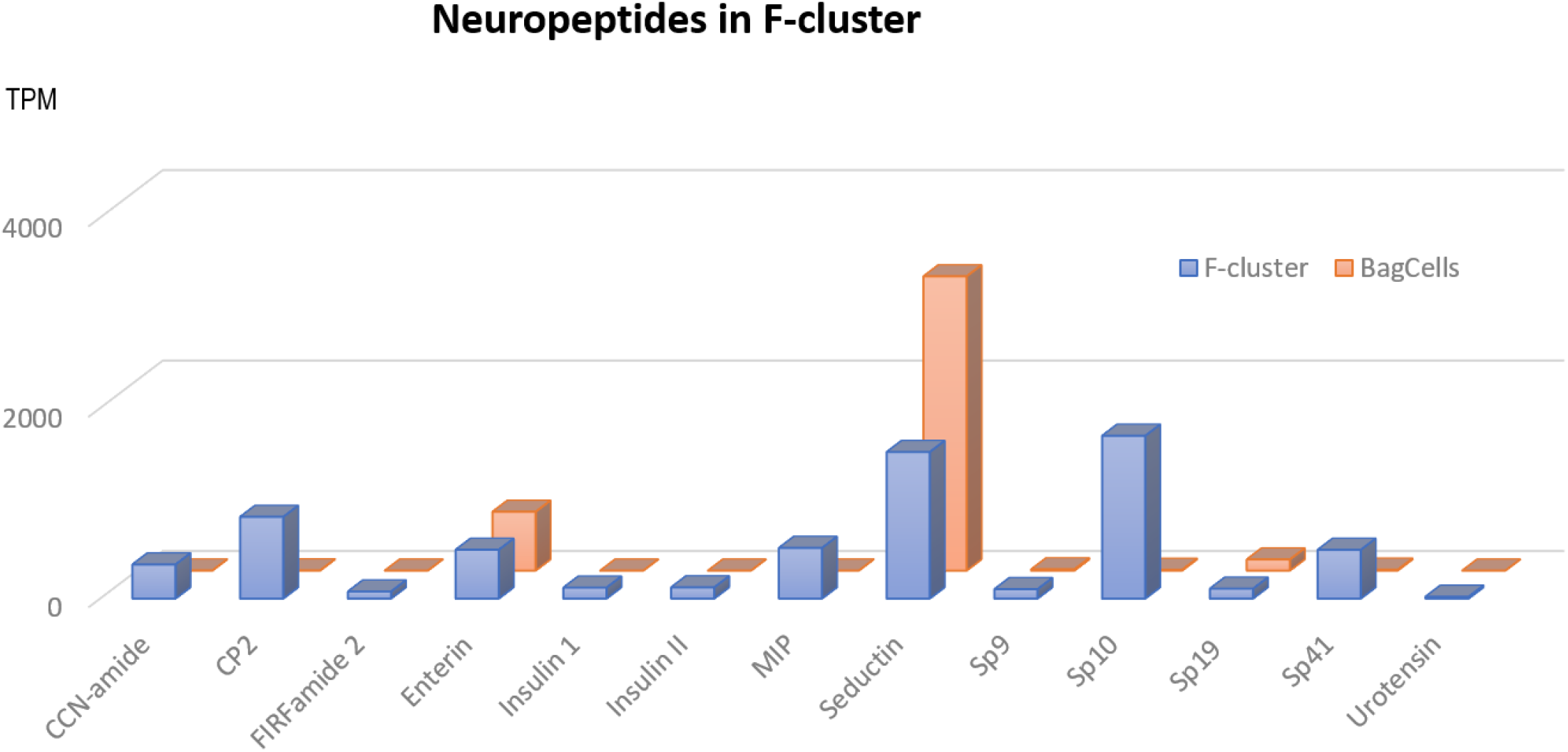
Expressions of selected neuropeptides in F-cluster and bag cells. (RNA-seq data). TPM - transcripts per million. See text for details.

**Fig. 7S.**
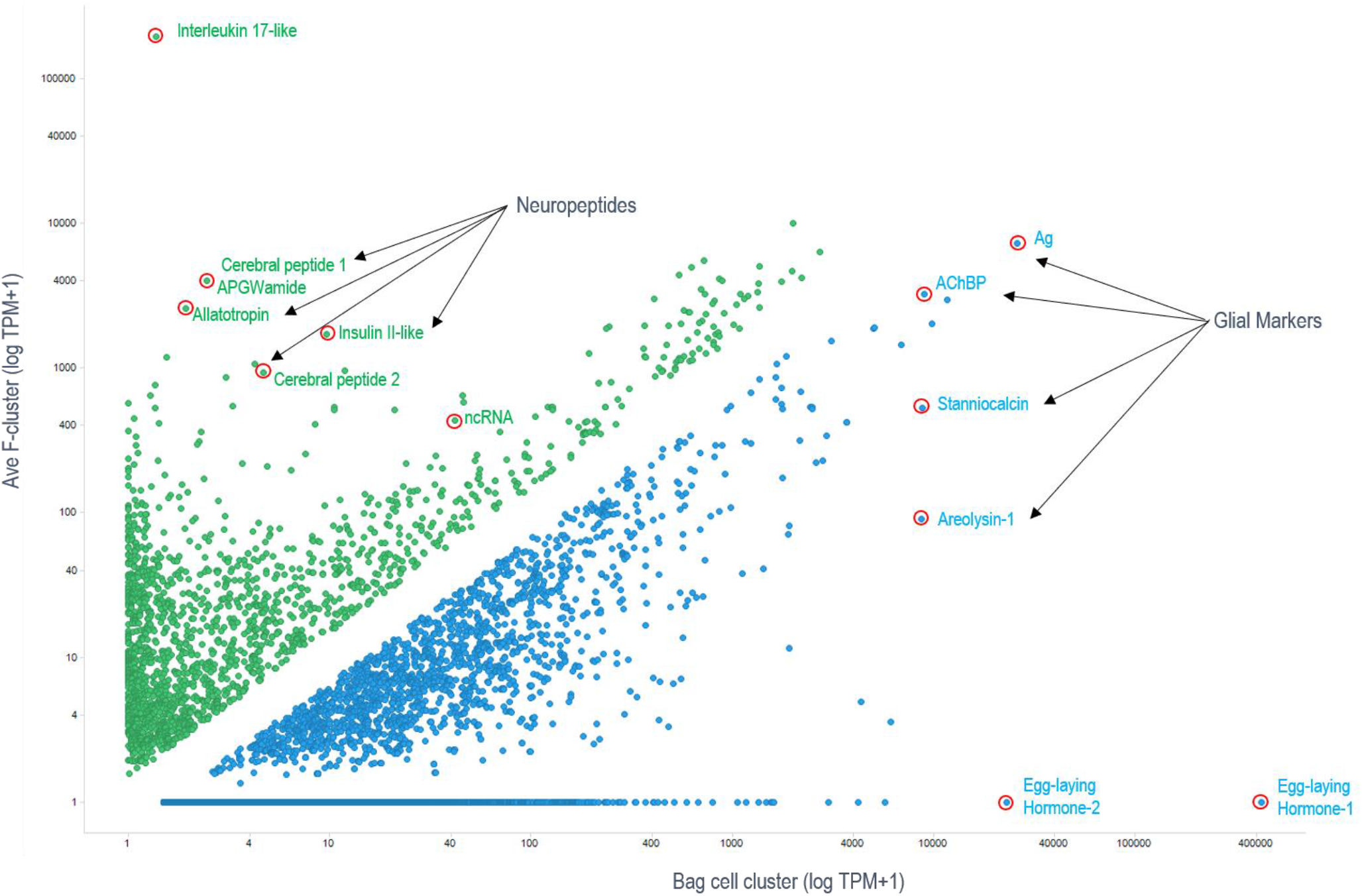
Illustrative. Examples of differentially expressed genes between F-cluster (green) and Bag cells (blue), RNA-seq data. TPM - transcripts per million. Of note, transcripts encoding interleukin 17-like and egg-laying hormones are the most abundant secretory products in F-cluster and bag cells, respectively. See text for details.

## Supplementary information 2

### Supplementary Methods & Sequences Descriptions

#### 1. Animals

*Aplysia californica* (60 - 100 g) were obtained from the National Resource for *Aplysia* at the University of Miami. Animals were anesthetized by injection of 60% (volume/body weight) isotonic MgCl_2_ (337 mM) prior to the removal of the CNS. The ganglia were then pinned to a sylgard dish in artificial seawater (ASW: 460 mM NaCl, 10 mM KCl, 55 mM MgCl_2_, 11 mM CaCl_2_, 10 mM HEPES, pH=7.6) and the cells were exposed by the mechanical removal of the overlying sheath with fine forceps.

#### 2. Molecular methods

##### 2.1 Cloning of Aplysia AcP2X4 receptors

The original sequences were obtained from *Aplysia* RNA-seq profiling(41,44,45,56). Details for RNA extraction and cDNA library construction have been described(41,44,56,57).

Both 5’ and 3’ RACE were performed to obtain the full-length coding sequence. Full-length CDS sequence for *Ac*P2X (GenBank# NP_001191558.1) was obtained using terminal primers: 5’-ATGGCTCCACCACAAGTCATGAAG-3’ and 5’-AAGCATCAAGGTGCGGCTCCTCCATCAC-3’. Amplified PCR product was cloned into pCR4-TOPO (Cat#K4575-01, LifeTechnologies). Four clones were isolated and sequenced together with a splice variant, *Ac*P2X_b_ (GenBank# NP_001191559.1), and the full-length CDS for this isoform also cloned and sequenced. Since there appears to be only one P2X gene in the *Aplysia* genome (GCF_000002075.1), we will designate the predicted protein as *Ac*P2X.

##### 2.2 Probe generation

The original sequences were obtained from *Aplysia* RNA-seq profiling(41,44,45,56). We used the same protocols for whole-mount *in situ* hybridization as reported elsewhere(58,59) We previously described for whole-mount *in situ* hybridization elsewhere(58,59) with a specific probe for the validated *Ac*P2X localization. The two isoforms of *Ac*P2X vary by a 147 base deletion/insertion and would not be distinguishable by *in situ*. The antisense probe was generated by digestion of the *Ac*P2X plasmid with Not I (Cat#R0189s, New England Biolabs Inc.) then transcribed with T3 polymerase from the DIG RNA Labeling Kit (Cat#11175025910, Roche Diagnostics). The control sense probe was produced by the same protocol but used Pme1 (Cat# R0560s, New England Biolabs Inc.) for digestion and T7 polymerase for transcription.

Expression of *Ac*P2X was investigated in central ganglia of 8 experimental and 2 control CNS preparations; additional controls were reported elsewhere(44,59). Control *in situ* hybridization experiments with full length ‘sense’ probes revealed no specific or selective staining in the CNS under identical conditions and labeling protocols. Images were acquired with a Nikon Coolpix 4500 digital camera mounted on an upright Nikon Optiphot-2 microscope.

##### 2.3. RNA and Oocyte preparation

RNA was transcribed from full-length cDNAs of *Ac*P2X and *Ac*P2Xb subunits using the T7 mMessage *in vitro* transcription kit (Ambion). The amount of purified, transcribed RNA was estimated on a Bioanalyzer (Agilent). Surgically removed stage V and VI oocytes from *Xenopus laevis* were injected with a total of 50ng transcribed RNA (46nL total volume) and incubated at 17°C for three-five days in ND96 medium (96mM NaCl, 2mM KCl, 1mM MgCl_2_, 1.8mM CaCl_2_, and 5 mM HEPES, pH=7.4) supplemented with 2.5 mM sodium pyruvate, 100 units/mL penicillin(Sigma), 100 μg/mL streptomycin(Sigma), and 5% horse serum(Sigma).

#### 3. Sequence analysis and phylogeny

Sequences were obtained through BLAST search across both Metazoans and non-metazoan groups and aligned using the MUSCLE or ClustalX ver. 2.1 programs with default parameters. Protein domains and motifs were obtained from Prosite(71) and SMART(72) databases. A maximum likelihood (ML) tree with the best-fit model (LG+G) was constructed using MEGA X(73) and 10,000 iterations.

Expression levels of transcripts were calculated using the normalization method for RNA-seq called Transcripts Per Million (TPM) as described in materials and methods. The projects used in this analysis are the following (**Table 1S**, below):

**Table 1S.** RNA-seq samples for *Aplysia californica*

| **SRA study SRP001185 (Development)** | **<u>SRA Run</u>** |
| --- | --- |
| Early Cleavage | SRX268443 |
| Gastrulation | SRX268442 |
| 3 Day | SRX268440 |
| 4 Day early trochophora | SRX268437 |
| 9 Day veliger | SRX268438 |
| 12 Day hatching veliger | SRX268439 |
| **BioProject PRJNA77701 (Tissue)** |  |
| Chemoreceptive tissue | SRX109676 |
| Heart | SRX109675 |
| Hepatopancreas | SRX109682 |
| Gills | SRX109674 |
| Salivary Glands | SRX109678 |
| CNS | SRX109680 |
| Muscle | SRX109681 |
| Other digestive parts | SRX109679 |
| Ovotestis | SRX109677 |
| Hermaphroditic glands and other reproductive organs | SRX109683 |

#### 4. The topology of predicted AcP2X4-1 and AcP2X4-2

##### 4.1 Secondary Structure

The deduced amino acid sequence for *Ac*P2X and *Ac*P2X_b_ contains two predicted transmembrane regions (TM1 and TM2), a large extracellular loop, and the N and C termini in the cytoplasmic face(33) supplement (**Fig.1S**, below). Both isoforms contain the ten conserved cysteine residues proposed to form sulfhydryl bonds(74). Also present in both subunits is the K-X-K-G motif on the N-terminal shown to be critical for ATP binding(75,76).

Both *Ac*P2X and *Ac*P2X_b_ contain five N-glycosylation sites fond mainly in the putative extracellular loop region, *Ac*P2X contains an additional site in the extracellular loop. Secondary motifs were determined with Prosite(71). Both predicted isoforms contain seven putative Protein kinase C (PKC) phosphorylation sites; each contains one site with a different location due to the splice region. Of the PKC sites, both *Aplysia* subunits contain the critical site involved in desensitization along with the conserved positively charged amino acid residues also involved in desensitization(75,76). *Ac*P2X and *Ac*P2Xb contain four Casein kinase II phosphorylation (CK2) sites, with *Ac*P2X having an additional two sites in its extracellular loop.

##### 3.2 Genomic Structure

The genomic organization of Human P2X4, NP_002551.2; Mouse P2X4, NP_035156.2, *Aplysia californica* P2X4-1, NP_001191558.1; *Capitella capitata*, ELU13670.1; *Daphnia pulex*, EFX89098.1; and *Nematostella vectensis*, XP_032239188.1 are compared. Human P2X4 and mouse P2X4 have 12 coding exons. *Aplysia* P2X receptor genes all have 11 coding exons in which the 9 and 10 exons compared to vertebrates are fused; however, they still generate a conserved exon/intron boundary. The asterisks indicate two contigs in the *Aplysia* genome assembly. All the exon/introns boundaries conserved and have compatible gene size. The two other invertebrate P2X-like receptor sequences from *Daphnia* and *Nematostella* only have 8 coding exons with 4 exons being fused. However, the exon/intron boundaries of those present in their genomes are still conserved. The exons are color-coded with the appropriate protein domain as well as critical amino acids being indicated.

